# PEtab Select: specification standard and supporting software for automated model selection

**DOI:** 10.1101/2025.05.12.653312

**Authors:** Dilan Pathirana, Frank T. Bergmann, Domagoj Doresic, Polina Lakrisenko, Sebastian Persson, Niklas Neubrand, Jens Timmer, Clemens Kreutz, Harald Binder, Marija Cvijovic, Daniel Weindl, Jan Hasenauer, the PEtab Select community

## Abstract

A central question in mathematical modeling of biological systems is determining which processes are most relevant and how they can be described. There are often competing hypotheses, which yield different models. Model comparison requires parameter optimization and sampling methods. Yet, standards for the specification of model selection problems and the swift evaluation of a broad spectrum of approaches are not available.

PEtab Select addresses this challenge by providing a concise, standardized specification of model selection and its associated calibration problems through a new file format standard and software package. The standard facilitates the compact representation of even very large model selection problems; in one example, billions of model alternatives. PEtab Select builds on the PEtab standard for the specification of parameter estimation problems, and enables the use of state-of-the-art modelling and calibration workflows utilizing COPASI, Data2Dynamics, PEtab.jl, and pyPESTO. PEtab Select supports common model selection criteria (e.g., Akaike and Bayesian information criteria) and can be easily extended to use others. To ensure flexibility, PEtab Select implements several model space exploration approaches, including basic brute-force, forward, and backward selection, and also advanced, flexible selection methods.

PEtab Select introduces the first standardization of model selection tasks, filling a critical gap in existing computational pipelines. It constitutes an essential contribution to FAIR research software in systems biology by promoting interoperability and reusability in model selection.

**Author summary:** Model selection is a crucial step in mathematical modeling, guiding the choice of components to include in a model. PEtab Select automates this process across diverse modeling frameworks and programming languages via (1) a new interoperable, language-agnostic standard for specifying large-scale model selection problems, and (2) a comprehensive software package that implements these selection methods. Developed through a community effort, PEtab Select has been integrated into multiple modeling frameworks and is accessible to users of COPASI, Julia, MATLAB, and Python.

## Introduction

In systems biology, mechanistic models explain and predict dynamic processes, guide experimental design, and support hypothesis testing [1, 2]. Because models encode biological assumptions in mathematical form, uncertainty arises both from incomplete hypotheses and from imperfect mathematical implementations. Model selection helps resolve these ambiguities by quantifying how well competing model formulations explain observed data [3].

The model-building cycle is increasingly supported by community standards. Models can be encoded in SBML [4], while simulations and parameterization workflows can be described using SED-ML [5] and PEtab [6]. Indeed, PEtab already allows one to specify measurement data, experimental conditions, observation models, and parameters, enabling interoperable calibration and analysis pipelines across tools and programming languages. Building on this, several software tools also offer model selection capabilities, including information criteria (AIC, AICc, BIC) [7–9], Bayes factors [10], likelihood ratio tests [11], and cross-validation [12]. Related functionality includes feature selection and model reduction to improve identifiability [13–18], as well as multi-model inference [19]. Widely used and versatile tools include Data2Dynamics (D2D) [20], dMod [21], and pyPESTO [22]. However, while model selection is possible using these tools, model selection problems are typically specified in tool-specific or ad hoc ways, limiting reproducibility and interoperability. Furthermore, support for different model selection approaches remains fragmented.

To address this gap, we introduce the PEtab Select standard for encoding model selection problems in systems biology. Developed in a community effort, the standard extends SBML [4] and PEtab [6] to provide a compact, interoperable description of model spaces and associated calibration problems, enabling large-scale comparative analyses without requiring new toolchains. Alongside the standard, we provide a reference PEtab Select software package that integrates into established calibration workflows. The package interfaces with PEtab-compatible tools such as Data2Dynamics [20], PEtab.jl [23], pyPESTO [22], and COPASI [24] via BasiCO [25]. It supports multiple exploration strategies—including brute force, forward, backward, and FAMoS [26]—while delegating model calibration to the selected backend tool. The following sections describe the standard and package and how they combine with existing workflows to streamline model selection (Fig. 1).

**Fig 1.**
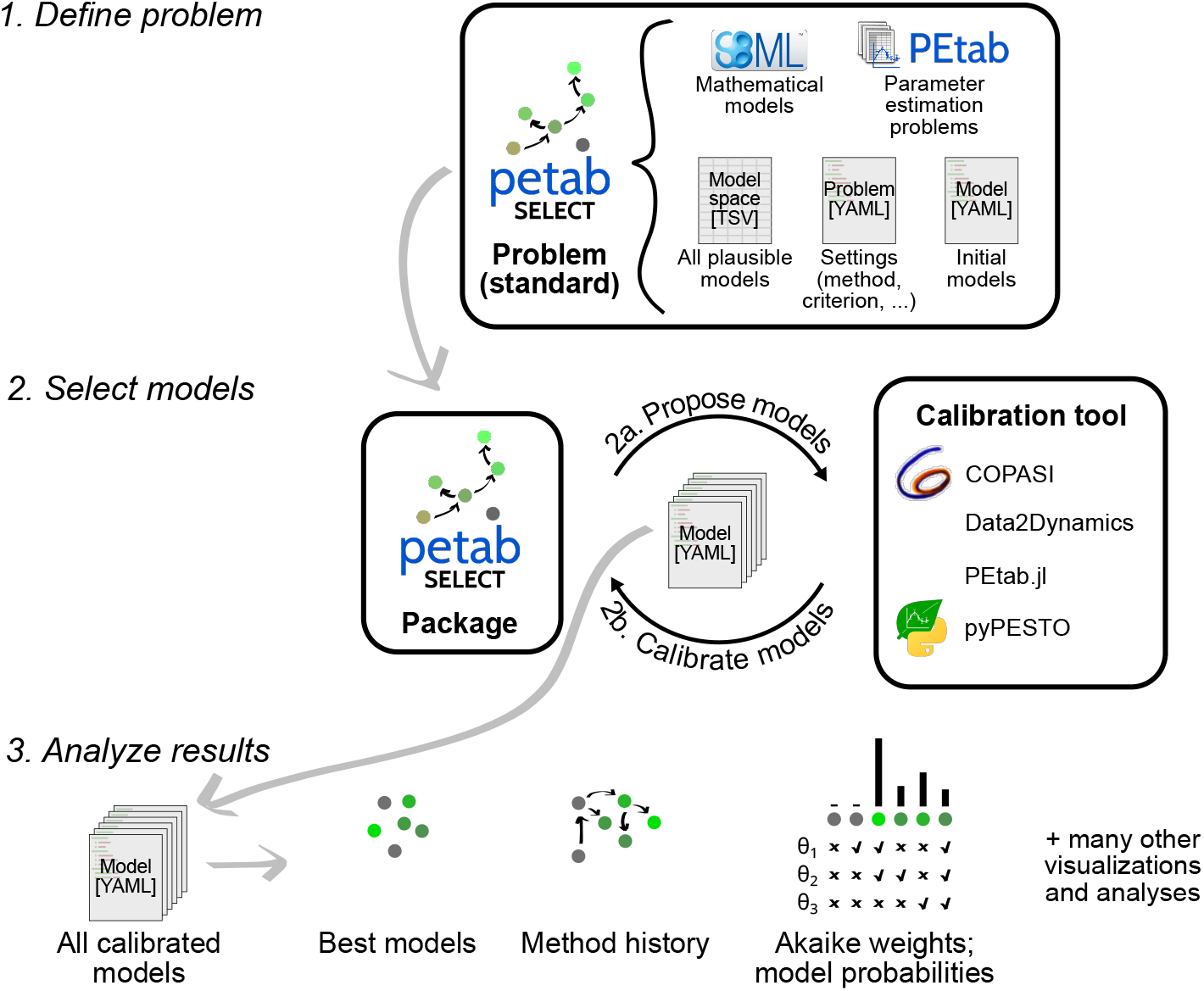
Model selection workflow with PEtab Select. (1) The user defines the model selection **problem**, including the model space and the mathematical description of the set of possible process components, in the **standard**. (2) The user runs model selection via a **calibration tool** by sending the PEtab Select problem to the PEtab Select **package**, which initiates iterations of model selection. (2a) The package provides model candidates to the calibration tool based on the problem specification and the available calibration results. (2b) The calibration tool calibrates the proposed candidate models and sends the calibration results to the package. Iterations of (2a) and (2b) continue until the package sends a termination signal to indicate the end of model selection. The results are provided in the PEtab Select model format and include all calibrated models with their calibration results. (3) The package can be used to perform subsequent analyses and visualizations of the results.

### Design and implementation

#### Scope

PEtab Select is designed to make model selection tasks portable across tools by separating (i) a standardized, tool-agnostic description of the model selection problem from (ii) the numerical calibration engine that evaluates candidate models. Concretely, PEtab Select provides (1) a concise specification (standard) that describes a model selection task in terms of candidate model definitions and associated calibration problems, (2) a reference software package that orchestrates model space exploration and model comparison, and (3) standardized output that records calibrated models and selection results for downstream analysis (Fig. 1). Calibration itself is performed by an external PEtab-compatible tool, allowing users to retain their preferred solvers, optimizers, and workflow-specific features.

#### The PEtab Select standard

The standard extends **PEtab** using the same TSV/YAML-based structure while adding model-selection-specific configuration. It preserves the PEtab description of parameter estimation problems (parameters, conditions, observables, and measurements), and adds a layer for defining the candidate model set and selection settings (e.g., exploration method and selection criterion). This ensures existing PEtab workflows remain valid, while enabling large and complex model selection problems to be specified compactly and exchanged between tools.

##### Model space (TSV)

The model space specifies which candidate models are admissible for selection. In PEtab Select, model spaces are represented as a combination of (i) explicitly listed single models and (ii) *superset models* from which multiple candidates are derived by varying designated components through a table of plausible parameter values. This permits compact specifications of large combinatorial spaces. In the current scope, candidate models are derived by parameter-level switches and value assignments (e.g., “estimate” versus fixed values), which covers common use cases such as including/excluding processes, reactions, or effects via parameterization.

An example of defining a model space based on two superset models is given in Fig. 2. Each superset model has an associated mathematical model and parameter estimation problem specified in the `model_subspace_petab_yaml` column. The set of submodels is constructed by customizing the superset model with all combinations of *plausible parameter values* in the additional table columns; for this specific example, subspaces `supersetA_models` and `supersetB_models` encode 16 (Fig. 2d) and 8 models, respectively. Note that for the example superset models in Fig. 2b and c, processes are turned “off” by setting parameters to 0, which is not always the case. For example, a fold-change effect parameter might be turned off by setting it to 1.

**Fig 2.**
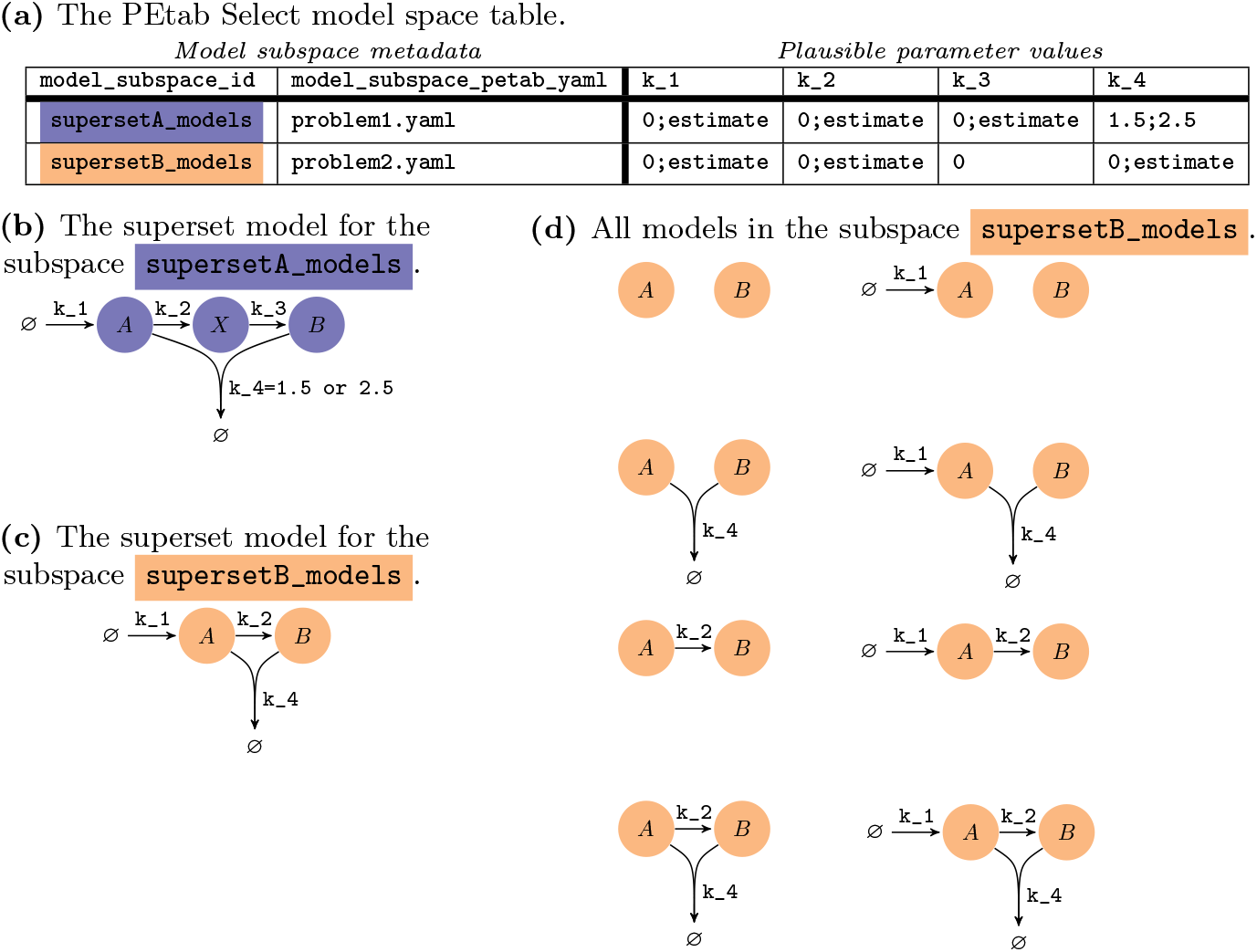
Example model space. (a) The model space table encodes model subspaces as rows. Each model subspace has an ID and an associated superset PEtab problem, which includes the mathematical model. This superset problem is customized into single models according to the remaining columns, which contain semicolon-delimited lists of values that parameters (column headers) can take in the subspace. (b,c) Each subspace is associated with a superset model, where all possible parameters are estimated. (d) Based on a superset model, candidate models are encoded by turning (estimation of unknown) parameters on or off.

##### Model files (YAML)

Given mathematical models, parameter estimation problems, and the model space, the package can run model selection methods. As the methods require calibrated models, the package sends the uncalibrated models to a calibration tool, which calibrates the models and then sends the results back to the package (step 2 in Fig. 1). The standard model file format is used to transmit all individual model information at all steps of the workflow (Fig. 1). These files are generated automatically by the package, but can also be supplied manually, for example to specify initial models (as done in test case 0009 of the PEtab Select test suite).

##### Model selection problem file (YAML)

As in PEtab, a single YAML-based “problem” file bundles the complete model selection task so compatible tools can load it from one entry point. The problem file links the relevant inputs (e.g., model space and PEtab problems) and specifies the selection configuration, including the exploration method and the selection criterion.

##### The PEtab Select package

The PEtab Select package extends calibration workflows by providing an API that enables model selection while leaving model calibration to the backend tool. Users typically interact with the package indirectly through their calibration environment: they provide PEtab/PEtab Select files, initiate model selection, and obtain standardized output in the PEtab Select model format for analysis and visualization (Fig. 1).

##### Calibration tools

To support PEtab Select, a calibration tool must be able to calibrate PEtab problems and interface with the package API. The package is written in Python 3 and can be accessed via a Python API or via the command line for non-Python tools. A dedicated test suite supports validation of tool integrations. PEtab Select is currently supported by COPASI [24] via BasiCO [25], Data2Dynamics [1, 20], PEtab.jl [23], and pyPESTO [22].

#### Model selection methods

The PEtab Select package supports global and local model selection methods to explore a user-defined model space. In each iteration, the package proposes one or more candidate models, obtains their calibration results from the backend tool, evaluates the chosen selection criterion, and decides which model(s) to consider next. Bold terms below denote methods and features implemented in the package.

##### Global methods

For exhaustive exploration, the package provides a **brute-force** mode that calibrates and scores every model in the specified space. This mode is useful for small or moderate model spaces and for benchmarking, but becomes impractical for combinatorial spaces (e.g., when *n* components can be switched on/off, yielding up to 2^*n*^ candidates).

##### Local methods

For larger spaces, the package implements stepwise local exploration. The **forward** method iteratively adds components (or releases fixed parameters for estimation), the **backward** method removes them, and the **lateral** method performs swap moves that add and remove in the same step. These local methods operate from an initial model, which can be provided by the user or generated automatically by the package.

The package also supports **FAMoS** [26], a metaheuristic local strategy that switches between forward, backward, and lateral moves according to a customizable switching scheme. In addition, FAMoS supports a “**jump to most distant**” move to escape local neighborhoods [26].

##### Model selection criteria

All methods rely on a model selection criterion provided in the PEtab Select problem specification. The package supports common information criteria, including **AIC** [7], **AICc** [8], and **BIC** [9]. Criteria are used consistently across candidate models by evaluating the calibrated fit while accounting for model complexity, enabling automated ranking and decision-making during exploration [3].

##### Other features

The package provides several other features to enable flexible model selection. This includes automated summaries of the model selection run, custom or automated selection of initial models, limits on the number of models calibrated in any iteration, restarting of model selection runs with previous model selection results, and object-oriented programming such that other methods and selection criteria can be easily implemented.

#### FAIR data principles

The FAIR principles emphasize machine-actionable interoperability and reuse [27]. PEtab Select supports these goals by: using machine-readable TSV/YAML formats; defining a documented controlled vocabulary; providing schemas (JSON Schema) for YAML-based files; and, enabling end-to-end reproducibility of model selection tasks across tools by building on community standards (SBML and PEtab). Results are stored in the same standardized format, enabling consistent downstream analysis independent of the calibration backend (e.g., via package-provided visualization utilities).

## Results

### PEtab Select enables model selection in existing workflows, each with unique tool-specific features

To assess the interoperability of PEtab Select with different software suites, we tested its functionality in four PEtab-compatible tools: BasiCO (built on COPASI), Data2Dynamics, PEtab.jl, and pyPESTO. Each tool successfully passed the tests in the test suite, confirming that they can parse and execute model selection tasks defined in the PEtab Select standard. While the software tools have a broad spectrum of overlapping features, there are a few distinctive characteristics.

**BasiCO (COPASI)** provides the possibility of custom evaluation functions in place of the standard information criteria. Moreover, its programmable interface enables model-specific optimizer settings, so callbacks or post-processing steps can be incorporated into the model selection workflow. This flexibility allows users to adjust parameters on a per-model basis or to visualize results automatically after each calibration step.

**Data2Dynamics** supports transferring parameter estimates from one calibrated model to another, potentially reducing computation time when new models are introduced. Additionally, users can define custom calibration routines to adjust the workflow before, during, or after parameter fitting. For example, the number of multi-starts in an optimizer can be dynamically set based on the size of the parameter space, or diagnostic plots can be automatically generated to assess calibration quality.

**PEtab.jl** integrates with the Julia SciML ecosystem, tapping into highly efficient ODE solvers provided by `DifferentialEquations.jl` [28] and automatic differentiation routines [29]. These capabilities enable rapid prototyping of novel approaches and runtime-efficient parameter estimation [23].

**pyPESTO** supports ensemble-based approaches, for instance, using Akaike-weighted ensembles to combine the results from multiple candidate models. Its ready-to-use post-processing tools also facilitate structured output of calibration results and automated generation of diagnostic figures, streamlining the analysis of potentially large model spaces.

### PEtab Select enables reproducible model selection across workflows

To assess the comparability of results across workflows, we consider an exemplary model selection problem from the work of Blasi et al. (2016) [13] that studied the acetylation of the histone H4 protein. It features a superset model where dynamics are governed by a reaction network with 16 state variables and 32 motif-specific reactions (Fig. 3a). Each reaction describes the conversion of one biochemical species into another. The rate constant of each motif-specific reaction is either the shared basal parameter or a motif-specific parameter. The model selection problem is to determine which reactions are sufficiently explained by the shared basal parameter, and which reactions require a motif-specific parameter instead. This model selection problem differs from the original publication in that we omit the additional 11 “site-specific” models in the original publication, which add a negligible computational cost and are not amongst the best models. Our model selection problem of 32 plausible motif-specific parameters therefore involves a model space of 2^32^ ≈ 4.3 billion plausible models.

**Fig 3.**
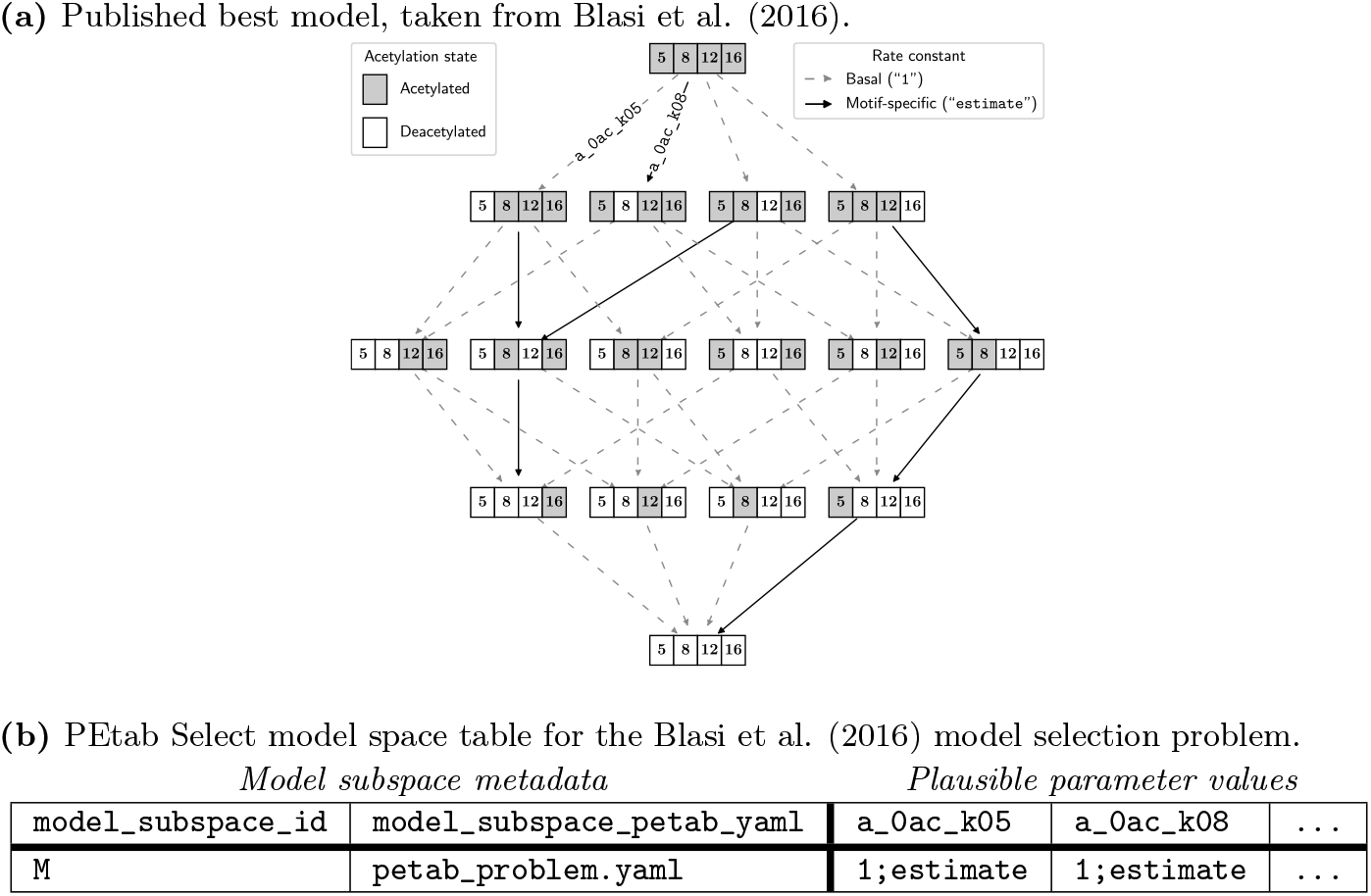
The Blasi et al. (2016) [13] model selection problem in PEtab Select. (a) The reaction network for the published best model. The histone chemical species can undergo acetylation reactions at four different positions (5, 8, 12, and 16). Nodes are acetylation states (motifs) of the histone, and arrows are acetylation reactions. The model selection problem involves two different possibilities for the rate constant of each reaction; it is either: the shared basal rate constant (the value is fixed to 1), or an estimated motif-specific rate constant (the value is estimated). The model selection problem is to identify the reactions with motif-specific rate constants, and the published best model contains seven (solid arrows). (b) There are 32 reactions, and therefore 32 independent hypotheses for motif-specific rate constants. Only two parameterized hypotheses (`a_0ac_k05` and `a_0ac_k08`) are shown here for brevity, with the remaining 30 parameters represented by the ellipses. This two-row, 34-column table encodes the full model space of ≈ 4.3 billion models.

The corresponding PEtab Select model space table is shown in Fig. 3b. The problem files and all technical details of the experiments in this section are provided in the Zenodo repository (see Availability and future directions).

In the original publication, all ≈ 4.3 billion models were calibrated then compared, and they found that a model with 7 motif-specific parameters is the best (Fig. 3a, and the Supplemental Information of [13], Table S1).

To solve our implementation of the model selection problem, we ran the PEtab Select implementation of the FAMoS method [26] 100 times, and each run was initialized at a randomly chosen model in the model space that had on average 16 motif-specific parameters (Fig. 4b). This first experiment with 100 FAMoS runs was to determine whether PEtab Select can find the same published best model, and was conducted using only pyPESTO as the calibration tool. The best model across the 100 searches matched the published best model, and the FAMoS algorithm found it three times in the 100 runs (Fig. 4c). In total, there were 83,125 (≈ 0.00194% of 2^32^; 58,858 unique) model calibrations across the 100 runs, which took 655.8 CPU hours.

**Fig 4.**
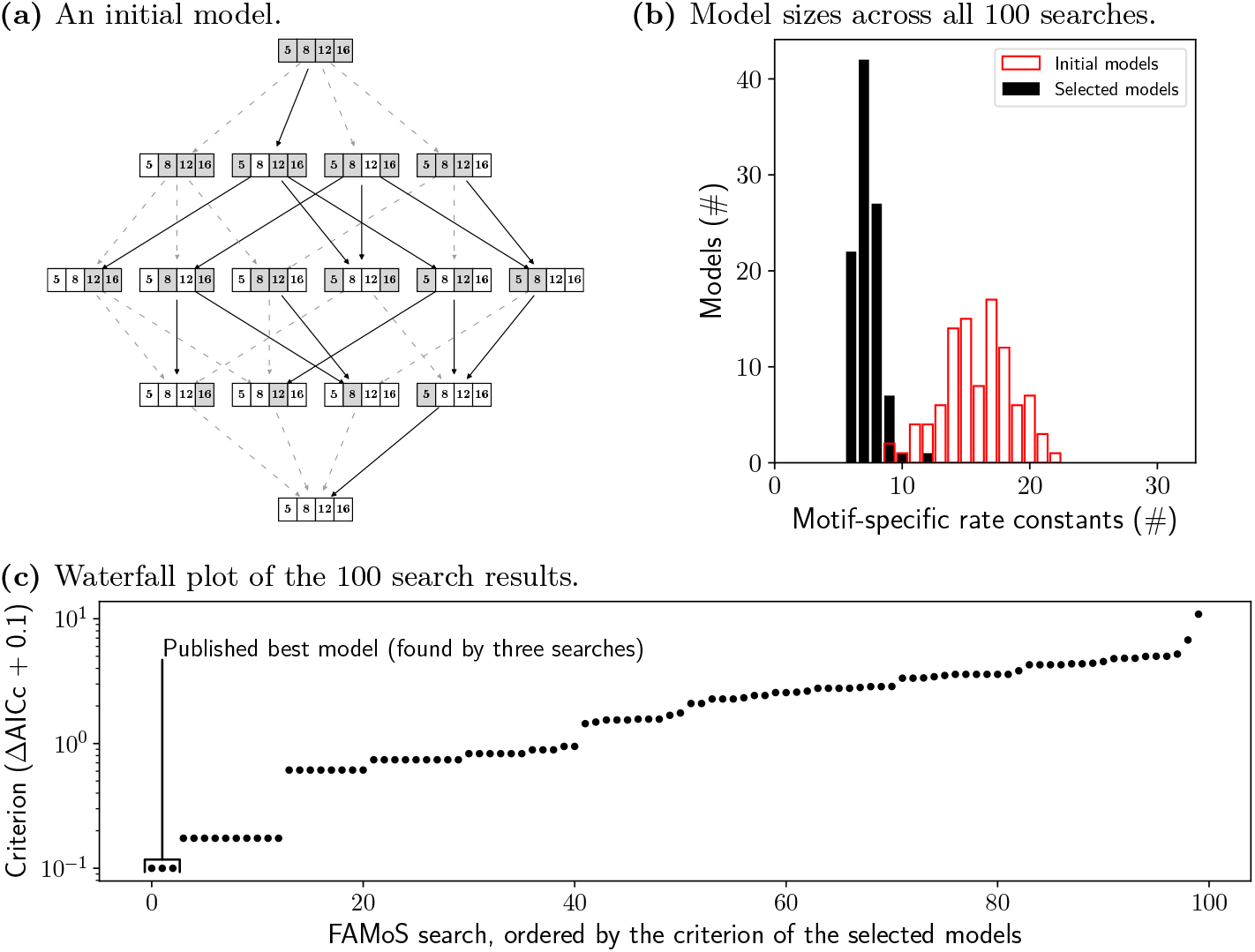
Multi-start FAMoS searches solve the Blasi et al. (2016) model selection problem. (a) The initial model that was used to assess reproducibility with all calibration tools, visualized as in Fig. 3a. (b) The sizes of models before and after model selection. The published best model had seven motif-specific rate constants. (c) The criterion of the selected model from each of the 100 FAMoS searches.

Extrapolating to the full model space, this saved 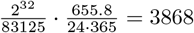 CPU years; hence, FAMoS rendered the standardized PEtab Select approach computationally feasible, meaning that tailoring the simulation and optimization routines to the problem, as was done in the original publication, is not necessary.

One of the three successful runs involved 339 model calibrations with pyPESTO, and its initial model is given in Fig. 4a. To assess reproducibility of model selection results generated with PEtab Select across all calibration tools, we repeated this FAMoS run with each tool.

All runs, i.e. with COPASI (BasiCO), Data2Dynamics and PEtab.jl, reproduced the same sequence of local search methods and calibrated similar sets of 339 models. Importantly, the same best model (Fig. 3a) was found with each of the calibration tools.

Model selection for the considered problem is inherently challenging. The search space is vast and populated by many candidate models whose information criterion values differ only within numerical precision, so minute floating-point variations render the ranking effectively non-deterministic across machines. Indeed, classical forward and backward selection mostly failed to arrive at the optimal solution. By contrast, the FAMoS algorithm [26] reproducibly converged on models with markedly better criterion scores. Although the reproduced single FAMoS search (with 339 model calibrations) takes slightly different trajectories through the model space on repeated executions within and between calibration tools (Fig. 5, calibration tool rows), each trajectory delivers the same overall improvement within simulator and optimizer tolerances (Fig. 5, row “Criterion (ΔAICc)”), with negligible differences (< 0.01) in criterion across calibration tools.

**Fig 5.**
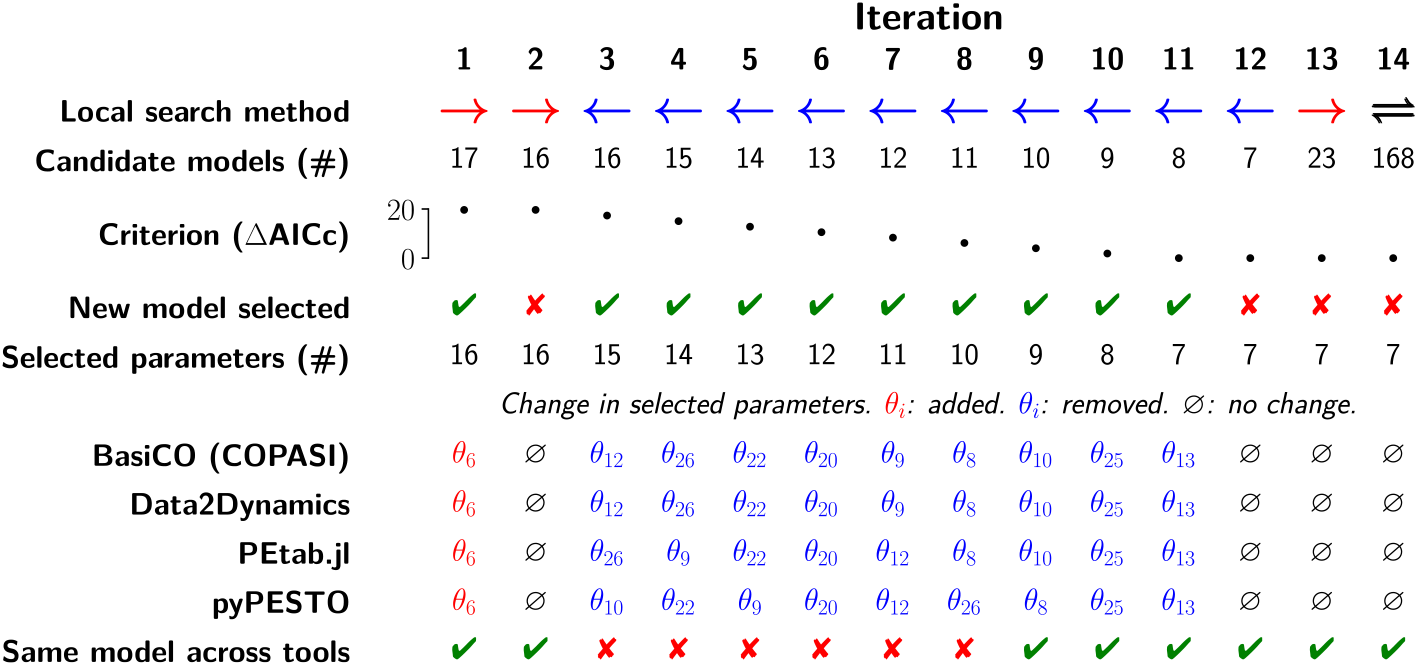
Reproducibility of model selection across calibration tools. Local search methods are the forward 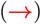, backward 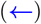, and lateral (**⇌**) methods. All calibration tools reproduced the same criterion value in each iteration, within 0.01 ΔAICc. The only difference between tools in each iteration is in the change in selected parameters. *θ*_*i*_ is the *i*th motif-specific parameter; see the Zenodo repository for details.

## Availability and future directions

Code, data, and specific package versions for reproducing the reported results is available at Zenodo (https://doi.org/10.5281/zenodo.17185040). Code for the PEtab Select package is maintained at GitHub (https://github.com/PEtab-dev/petab_select/), and documentation is hosted at ReadTheDocs (https://petab-select.readthedocs.io/).

There are several useful directions in which PEtab Select can be improved. Currently, PEtab Select only provides support for the SBML mathematical model format, as this is the only model format supported by PEtab. However, PEtab Select has no direct dependency on the model format. Hence, this is expected to improve with the next version of PEtab, which will support non-SBML model formats. Further implementation improvements include shifting calibration-tool specific features into the package directly, such that those features can be used with any calibration tool. One example would be the termination of a model selection iteration as soon as a better model is found.

Several opportunities for the improvement of model selection methodology were identified when working with the problem published by Blasi et al. (2016) [13]. On one hand, the analysis revealed that reproducibility could be boosted by developing tailored strategies for handling the case that multiple models are equivalent in best criterion value (up to numerical noise). If performing a backward search, one possibility is to branch the backward search into multiple backward searches, one per equivalent model. On the other hand, a further improvement of search algorithms would be beneficial, in particular for large model spaces, e.g., an automatic tuning of metaheuristics in the FAMoS method across runs.

In summary, PEtab Select establishes, for the first time, a unified standard and reference implementation for large-scale, reproducible model selection. By decoupling the description of a model selection task from the calibration engine that solves it, the framework enables researchers to tackle search spaces of billions of candidates in whichever environment—COPASI, Data2Dynamics, pyPESTO, PEtab.jl, or future tools—best suits their workflows. This interoperability not only accelerates day-to-day practice, but also lowers the barrier for methodological innovation, allowing new inference algorithms to be benchmarked immediately on realistic problems. We therefore expect PEtab Select to become a cornerstone for transparent, comparable, and scalable model selection across the life and physical sciences.

## Acknowledgments

The authors are grateful to the many people who have provided feedback and contributions to the standard, package, and other aspects of PEtab Select, including participants of the HARMONY and COMBINE conferences (https://co.mbine.org).

This work was supported by the TRA Modelling (University of Bonn) as part of the Excellence Strategy of the federal and state governments in Germany, the Deutsche Forschungsgemeinschaft (DFG, German Research Foundation) under Germany’s Excellence Strategy (project IDs 390685813 - EXC 2047 and 390873048 - EXC 2151) and through Metaflammation (project ID 432325352 – SFB 1454), the German Federal Ministry of Education and Research (BMBF) within the e:Med funding scheme (junior research alliance PeriNAA, grant no. 01ZX1916A), the Horizon Europe - ERC Consolidator Grant 2023 (grant agreement no. 101126146), the Federal Ministry of Education and Research (grant no. 031L0293C), and by the University of Bonn via the Schlegel professorship to JH. FTB was supported by LIBIS, and de.NBI, the German Network for Bioinformatics Infrastructure (W-de.NBI-016). MC and SP were supported by the Swedish Research Council (VR2017-05117 and VR2023-04319) and the Swedish Foundation for Strategic Research (FFL15-0238). The work of NN, JT, CK, and HB has been funded by the Deutsche Forschungsgemeinschaft (DFG, German Research Foundation) – Project-ID 499552394 – SFB 1597 Small Data.

## PEtab Select community

Severin Bang, Adrian Hauber, Svenja Kemmer, Daniel Lill, Lukas Refisch, Marcus Rosenblatt, Christian Tönsing, Jakob Vanhoefer, Franz-Georg Wieland.

